# Maladaptive cue-controlled cocaine-seeking habits promote increased relapse severity in rats

**DOI:** 10.1101/2021.02.01.429216

**Authors:** Maxime Fouyssac, Yolanda Peña-Oliver, Mickaёl Puaud, Nicole Lim, Chiara Giuliano, Barry J Everitt, David Belin

## Abstract

The inflexible pursuit of drug-seeking and great tendency to relapse that characterize addiction has been associated with the recruitment of the dorsolateral striatum-dependent habit system. However, the mechanisms by which maladaptive drug-seeking habits influence subsequent relapse are obscure. Here, we show that rats with a long history of cocaine-seeking, controlled by drug-paired cues and mediated by the habit system, show highly exacerbated drug-seeking at relapse that is not mediated by cocaine withdrawal. This heightened tendency to relapse is underpinned by transient engagement of the dorsomedial striatum goal-directed system and reflects emergent negative urgency resulting from the prevention of enacting the seeking habit during abstinence. These results reveal a novel mechanism underlying the pressure to relapse and indicate a target for preventing it.

**One Sentence Summary:** Instrumental deprivation triggers flexibility in the well-established cue-controlled cocaine-seeking behaviour.

## Main text

The initiation of drug use depends on the reinforcing properties of addictive drugs mediated by dopaminergic transmission in the nucleus accumbens (NAc) (*1*), which is also activated by motivationally relevant drug-paired conditioned stimuli (CSs) in recreational drug users (*2–4*).

However, in chronic drug users, or individuals with a diagnosed addiction (severe substance use disorder) (*5*), presentation of the same CSs also triggers the activity of, and dopamine release in, the dorsal striatum (DS) (*4, 6, 7*). Cue-provoked activation of the neural network converging onto the DS, often referred to as the habit system (*8, 9*), has been shown in individuals with an addiction to be a strong, if not the best, predictor of long-term relapse (*10*).

This transition in the neural locus of the impact of drug CSs from dopamine-dependent mechanisms in the NAc to encompass dopaminergic mechanisms in the dorsolateral striatum (DLS) over the course of a long history of drug use has been shown in humans (*2, 4, 6, 11*) and non-human primates (*12, 13*), as well as in the control over drug-seeking in rodents (*14–17*).

The psychological and behavioural significance of the functional engagement of the DLS during the development of drug addiction and the associated vulnerability to relapse are both complex and not fully understood. We have provided evidence that it reflects the development of drug-seeking habits in which individuals with an addiction rigidly engage to obtain and eventually take the drug (*15, 18–22*).

The development of DLS-dependent habitual control over drug-seeking has been suggested to be incompatible with the seemingly flexible and inventive behaviours that addicted individuals perform when actively engaged in drug-seeking activities, especially when obtaining the drug is made difficult by changes in context, such as scarcity of sources of money, unavailability of their regular drug supply or danger in the environment (*23*). This flexibility in the performance of drug-seeking is considered to reflect its goal-directedness, implying it is motivated by, and enacted following a neural representation of the value of the drug (*21, 24, 25*). However, flexibility in the performance of an ongoing drug-seeking behavioural sequence does not necessarily reflect the nature of the psychological mechanisms involved in its initiation (*26*).

Outside laboratory settings, this distinction between the psychological and neural basis of the initiation of drug seeking behaviour and its subsequent performance is especially important. In everyday life, drug-seeking occurs over prolonged periods, and the attendant delays to eventual drug use (*27–29*) are bridged by drug-associated CSs that capture attention (*30–32*) and, by acting as conditioned reinforcers (CRfs), maintain seeking responses while also playing a significant role in relapse (*33*).

Cue-controlled drug-seeking behaviour has been operationalized in second-order schedules of reinforcement (SOR) (*27, 34–36*) in which humans, non-human primates or rats will vigorously seek a drug over long time intervals (usually 15 min in rats), but only when their instrumental seeking responses are reinforced by the contingent presentation of a drug-paired conditioned stimulus (CS) (*37*).

At the neural systems level, the acquisition of cocaine-seeking under the conditioned reinforcing properties of cocaine-paired cues depends on a network involving the basolateral amygdala (BLA) (*38*), the nucleus NAc core (NAcC) (*39*), their functional interactions (*40*) mediated by direct BLA-NAcC projections (*41*) as well as the posterior dorsomedial striatum (pDMS) (*42*). When cocaine seeking is well-established through daily performance over several weeks, it becomes habitual and mediated by a network that involves the central amygdala (*43*) and dopamine-dependent mechanisms in the anterior DLS (aDLS) (*14, 44, 45*). However, the extent to which cocaine-seeking habits influence the high likelihood of relapse after abstinence seen in addicted individuals is not well understood.

In a first experiment (**Fig. S1)**, 47 rats were progressively trained to seek cocaine for prolonged periods under a SOR in which every 10^th^ instrumental seeking response was reinforced by the contingent presentation of the cocaine paired CS and cocaine was infused intravenously following the 10^th^ response after a 15-minute Fixed Interval had timed out (formally this schedule is denoted by FI15(FR10S)) (see **SOM** and **Fig. S2**). After at least two weeks of cue-controlled cocaine seeking, conditions previously shown to engage the aDLS in the control over this habitual seeking behaviour (*14, 22, 42*), a 3-day abstinence period was imposed on the rats prior to their reintroduction to the SA chamber in which they could again freely seek cocaine for 15 min in a drug-free state, precisely as they had been allowed to do in the pre-abstinence daily sessions (**Fig. S1, exp. 1a and Fig. S3**).

Compared to baseline levels of responding before these three days of imposed abstinence, SOR-trained rats (SOR rats) showed a marked (~200%) increase in seeking responses at this relapse test (**Fig. 1A**). This extremely high level of instrumental seeking was not observed in control rats that had a similar history of drug intake, but whose seeking responses had never been under the influence of cocaine-associated conditioned reinforcers (*46*) (henceforth referred to as FI15 control rats) (**Fig. 1A**).

**Figure 1:**
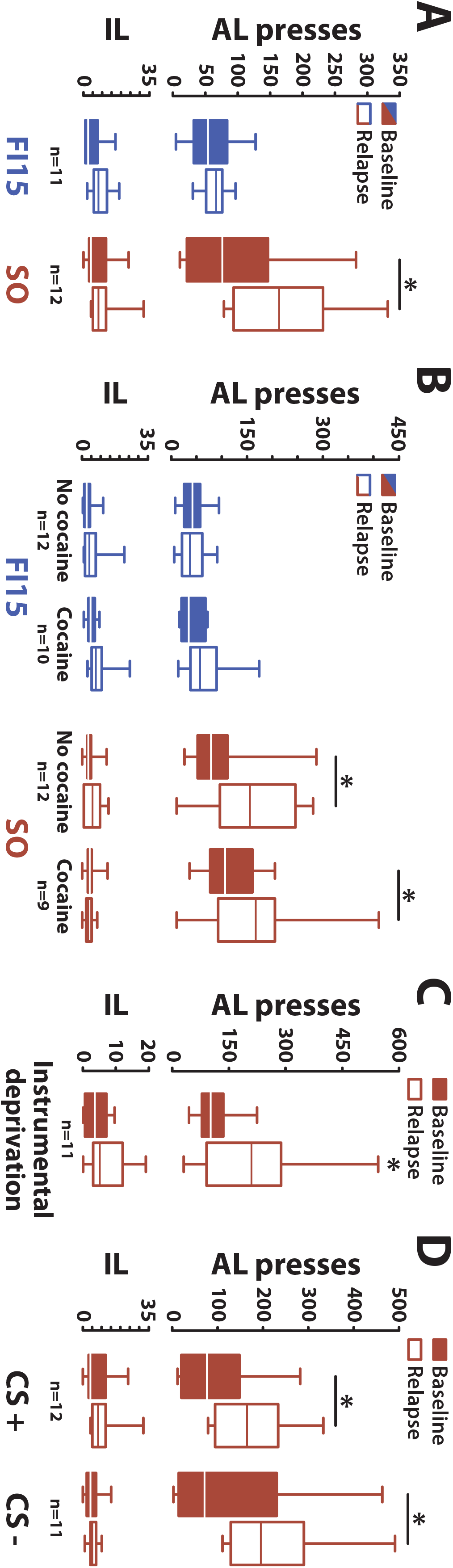
A prolonged history of cocaine seeking under the control of cocaine-associated conditioned reinforcers is associated with aberrantly enhanced drug-seeking as a consequence of instrumental deprivation. **A)** Following three days of forced abstinence after a prolonged history of cocaine-seeking under FI15 (FI15 rats, n=11) or SO schedules of reinforcement (SOR rats, n=12), only rats having developed cocaine-seeking habits under the SOR showed an aberrant and marked increase in seeking responses at relapse as compared to their baseline response levels [main effect of group: F_1,21_=9.43, p<.01, 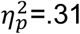; and lever × abstinence × group interaction: F_1,21_=4.43, p<.05, 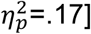. This instrumental deprivation effect was not due to a differential response rate between SOR- and FI15-trained rats as revealed by an analysis of covariance computing baseline level of responding as a covariate [main effect of group: F_1,39_=6.57, p<.02, 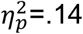, response rate: F_1,39_=1.94, p=0.17 and group × response rate interaction: F_1,39_<1]. The aberrant increase instrumental responding, or rebound, was shown not to be attributable to cocaine withdrawal (**B**), since a similar rebound by SOR, but not FI15, rats was seen even when daily cocaine infusions were given contingently at the dose rats would have self-administered during the 3 days of abstinence but, importantly, were not given the opportunity to seek the drug [main effect of group: F_1,39_=28.58, p<.001, 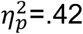; lever × abstinence × group interaction: F_1,39_=4.41, p<.05, 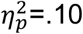 but no main effect of abstinence conditions, or any interactions involving abstinence conditions, all Fs<1]. **C)** Even when they were able to self-administer cocaine by making single taking responses every 15 minutes during the imposed ‘abstinence’ 3 days, SOR rats still showed the rebound increase in seeking responses at relapse [main effect of lever: F_1,10_=39.95, p<.001, 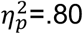; instrumental deprivation: F_1,10_=6.79, p<.05, 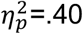; and instrumental deprivation × lever interaction: F_1,10_=6.54, p<.05, 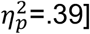. **D)** The instrumental deprivation effect only emerged following a long history of cocaine-seeking under the influence of cocaine-associated conditioned reinforcers, but its manifestation as a rebound in instrumental responding did not require the presence of these same CSs at relapse [lever × abstinence interaction: F_1,21_=8.40, p<.01, 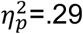; no main effect of CS presentation or any interactions involving CS presentation, all Fs<1]. *p≤.05

Since both SOR and FI15 control rats had the same quantitative cocaine exposure throughout their history and immediately prior to abstinence (**Fig. S2**), it was unlikely that the post-abstinence behavioural rebound shown only by the SOR rats was attributable to a drug withdrawal-associated motivational drive (negative reinforcement) (*47, 48*), but we nevertheless investigated this possibility.

Rats underwent another three days during which they were prevented from seeking cocaine, but during this imposed ‘abstinence’ period they now received daily, non-contingent intravenous administrations of the same number of cocaine infusions every 15 minutes as they would have obtained through their seeking behaviour (**Fig. S1, exp. 1b and Fig. S3**). When, on the fourth day, they were reintroduced to the SA chamber with the opportunity freely to seek cocaine for a 15 min drug-free period (**Fig. S3**), SOR rats showed the same 200% increase from baseline in instrumental responding at relapse while the FI15-control rats did not (**Fig. 1B**). This result suggests that the post-abstinence rebound in the vigour with which SOR, but not control, rats instrumentally respond was not the consequence of cocaine deprivation during imposed abstinence.

Non-contingent administration of cocaine does not, of course, recapitulate the complex psychological and physiological mechanisms engaged when self-administering the same dose of the drug (*49–51*). Hence, in an additional experiment **(Fig. S1, exp. 1c and Fig. S3**), rats were placed in the SA chamber and given the opportunity to take, but not to seek, cocaine by allowing them to make a single taking response for a cocaine infusion after each FI 15 mins of the session during what was otherwise three days of imposed abstinence, but they were not allowed to perform seeking responses. When rats were again given the opportunity on the 4^th^ day freely to seek cocaine, SOR rats showed the now characteristic 200% increase in responding compared to their baseline levels (**Fig. 1C**).

These results show that the increased responding at relapse shown specifically by SOR rats is independent of both deprivation of, or withdrawal from, cocaine as it occurred when rats were maintained on the same daily non-contingent doses of cocaine and even when allowed to self-administer it during the three imposed ‘abstinence’ days. Furthermore, the rebound effect did not depend on any differences in the initial acquisition of cocaine SA (**Fig. S2**) nor, as shown by analysis of covariance, on differences in the baseline rate of responding between SOR and FI15 control rats (**Fig. 1**).

The extremely high level of seeking behaviour shown specifically by SOR rats can only be attributed to the prevention of the opportunity to express their well-established, habitual seeking responses on abstinence days. It is more appropriately viewed as an instrumental deprivation effect.

An additional, intriguing feature of the aberrantly greater instrumental seeking behaviour after imposed abstinence is that it was not dependent on the contingent presentation of cocaine-associated CSs in the relapse test (**Fig. S1, exp. 1d**). Omission of the CRf at the relapse test after three days of imposed abstinence did not result in lower levels of seeking by SOR rats (**Fig. 1D**), even though it had greatly increased their seeking behaviour under baseline conditions. We therefore conclude that it is the history of drug-seeking under the control of drug-associated CSs, which we have previously suggested to result in the formation of incentive habits (*21, 52*), but not the presence of these same response-contingent CSs at test, that drives the instrumental deprivation effect. This effect clearly differs from the incubation of craving, which reflects a marked increase in responding for a drug-associated CS under extinction conditions that develops in rats after long periods of abstinence (*33*).

Even though the SOR and FI15 control rats in these experiments had a prolonged history of cocaine-seeking prior to abstinence, they had not experienced the extended access conditions that result in escalated cocaine intake (**Fig. S1, exp. 2**) indicating loss of control over intake (*53*). We therefore investigated whether the instrumental deprivation effect may be qualitatively or quantitatively influenced by a previous history of loss of control over cocaine intake. A further cohort of 24 rats acquired cocaine SA under FR1 over eight 1-hour daily sessions and then underwent either twelve additional 1-hour daily sessions (short access, ShA) or 12 6-hour extended access sessions (long access, LgA), until a robust escalation of intake was observed (**Fig. S4 A-C**). Rats were then trained and maintained for two weeks to seek cocaine under a SOR, as in the previous experiment (**Fig. S4 D-E**). ShA and LgA rats did not differ in their levels of responding under a SOR, nor in the magnitude of the instrumental deprivation effect at relapse following three days of imposed abstinence (**Fig. S4 F**), which, again, was shown to be independent of the presentation of the cocaine-associated CRf at test.

The motivational mechanisms involved in the rebound in instrumental responding following instrumental deprivation hence seem unlikely to depend, for example, on adaptations, in the stress system that are recruited following extended exposure to, or withdrawal from, the drug (*48*), but emerge instead as a consequence of a prolonged history of drug seeking under the influence of the conditioned reinforcing properties of response-produced cocaine-paired Pavlovian cues (*21, 52, 54*).

Taken together these data suggest that several weeks of drug seeking under conditioned reinforcement results in an urge for action (*55*) that emerges when animals are deprived of the opportunity to express this well-consolidated, or ingrained, cocaine seeking habit. This urge to act (*55*) following instrumental deprivation may resemble the motor urges characteristic of impulsive/compulsive disorders such as Tourette syndrome (*56*) or compulsive eating (*57*), that have been suggested to be a manifestation of the negative urgency (*58–60*) that can precede and mediate relapse (*59, 61*).

We hypothesized that the post-instrumental deprivation rebound of responding at relapse reflects a behavioural response driven by an urge to act that is goal-directed, but the goal is not the drug, but to respond and hence alleviate the negative state associated with the urge (*59*). We further hypothesized that rather than being mediated by the aDLS-dependent habit system on which well-established cue-controlled cocaine seeking has been consistently shown to rely (*14, 41, 42, 44, 62*), the rebound would instead depend on re-engagement of the pDMS goal-directed system (*63, 64*).

To initially explore the possible neural locus of control of this aberrant rebound of responding at relapse, mRNA levels of the neuronal activity marker c-Fos were measured throughout the DS (*65*) using in situ hybridization after a test session following either baseline performance or instrumental deprivation.

After a prolonged history of cue-controlled cocaine-seeking, three days of imposed abstinence from cocaine and the ability to seek it, SOR rats again showed the characteristic rebound of instrumental responding at relapse as compared to their non-abstinent controls (**Fig. 2A-left**). Neurally, this instrumental rebound at relapse was associated with an increase in c-Fos mRNA levels in the pDMS, whereas they were unchanged in the aDLS (**Fig. 2A-right**).

**Figure 2:**
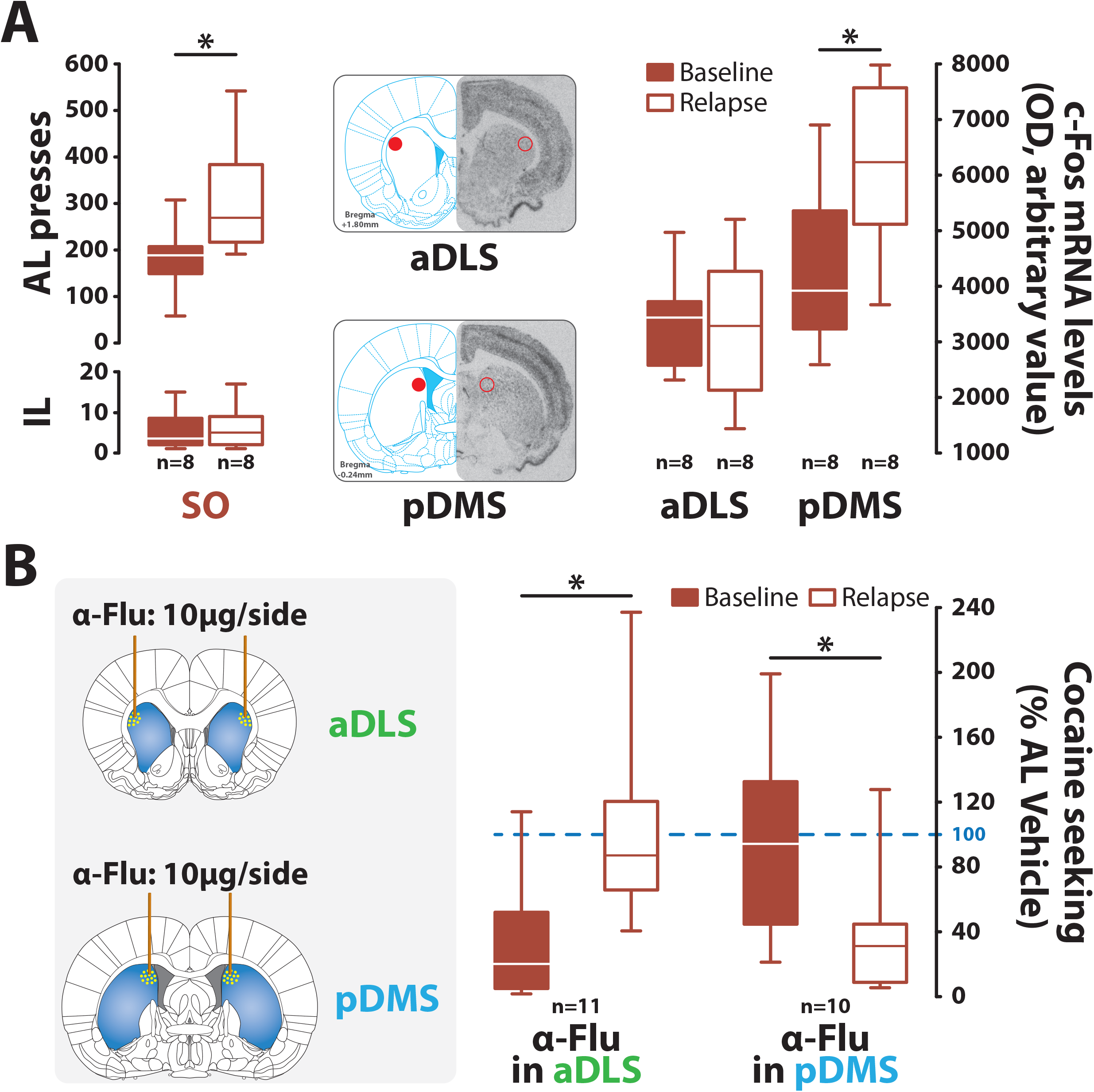
The exacerbated tendency to express drug-seeking responses following instrumental deprivation depends on the transient re-engagement of the goal-directed system. **A)** Following three days of abstinence after a prolonged history of cocaine-seeking under a second order schedule of reinforcement (SOR rats, n=8), rats showed the characteristic aberrant engagement in instrumental responding at relapse as compared to their non-abstinent controls (SOR rats, n=8) [main effect of abstinence: F_1,14_=6.20, p≤.05, 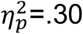, lever: F_1,14_=79.38, p<.001, 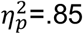 and abstinence × lever interaction: F_1,14_=5.81, p≤.05, 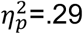; left panel]. The rebound of seeking behaviour after instrumental deprivation was associated with an increase in c-Fos mRNA levels in the pDMS [F_1,14_=6.06, p≤.05, 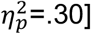 but not in the aDLS [F_1,14_<1]. This observation suggested that the instrumental deprivation effect was associated with a functional engagement of the pDMS in the control over behaviour. **B)** A causal investigation of the reliance of instrumental seeking responses at baseline and following instrumental deprivation on aDLS- or pDMS-dopamine dependent mechanisms was subsequently undertaken on an additional cohort of eleven rats by measuring the sensitivity of seeking responses to intracerebral infusions of the dopamine receptor antagonist α-flupenthixol infused into each striatal domain (10 μg/side) (*14, 22, 42, 66*). Following a long history of cocaine-seeking under a SOR instrumental seeking responses were decreased by 50% by intra aDLS α-flupenthixol infusions relative vehicle infusions while intra pDMS receptor blockade had no effect under these baseline conditions. In marked contrast, following three days of imposed abstinence, intra-pDMS infusions of α-flupenthixol resulted in a decrease of over 60% of the rebound increase in seeking, whereas intra aDLS α-flupenthixol infusions were now without effect [main abstinence × locus interaction F_1,19_=16.33, p<.001, 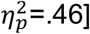. *p≤.05

This observation, consistent with our prediction, suggests that the post-instrumental deprivation rebound of responding is associated with a transient functional engagement of the pDMS in the control over behaviour at a time when drug seeking at baseline prior to abstinence is mediated by aDLS dopamine-dependent mechanisms (*14, 43*).

We causally tested this hypothesis on two additional cohorts of 24 rats each implanted with cannulae targeting either the aDLS or the pDMS (**Fig. S1** and **S5**) and trained to seek cocaine under SOR for at least three weeks (**Fig. S1** and **S6**), conditions known to result in engagement aDLS-dopamine dependent control over behaviour (*14, 42, 43*). The effects on instrumental responding of infusing the DA receptor antagonist α-flupenthixol (*3, 37, 66*) into the pDMS or the aDLS prior to testing were investigated following instrumental deprivation (i.e., three days of imposed abstinence), or under baseline (no abstinence) SOR conditions (**Fig. S1** and **S6**).

As expected from our earlier results (*14, 42, 43*), well established cue-controlled cocaine-seeking responses at baseline were decreased by over 50% following intra-aDLS α-flupenthixol infusions relative to vehicle infusions, whereas responding was completely unaffected by intra pDMS dopamine receptor blockade (**Fig. 2B**). In marked contrast, following three days of imposed abstinence, intra aDLS α-flupenthixol infusions had no effect on the rebound in instrumental responding following instrumental deprivation, which was instead significantly decreased by over 60% by dopamine receptor blockade in the pDMS (**Fig. 2B**).

This double dissociation shows that in the same individual, at a time when habitual cue-controlled drug-seeking is under aDLS dopamine-dependent control (*14, 42*), the maladaptive engagement of cocaine seeking following abstinence-induced instrumental deprivation is associated with a functional switch in the locus of control over behaviour to the pDMS goal-directed system (*42, 64, 67*). Importantly, this engagement of the pDMS following instrumental deprivation occurs on a background of maintained activation of the aDLS, as measured by c-fos mRNA levels, suggesting alteration in the functional balance between the goal-directed and habit systems in this new context (*65, 68–70*).

The transient re-engagement of pDMS control over behaviour at relapse is not attributable to the engagement of a cocaine-withdrawal motivational state, nor even to loss of the opportunity to self-administer cocaine, since maintaining rats on their usual dosing of cocaine during the three days of imposed abstinence did not prevent the rebound effect.

Instead, we propose that it reveals the recruitment of goal-directed control over behaviour when individuals have been deprived of the opportunity to perform their drug seeking responses, but that the goal is no longer the drug, but the alleviation of an urge to perform their seeking responses that themselves have acquired motivational value.

This rebound in instrumental responding and the underlying switch to pDMS control exacerbates the severity of relapse, but it cannot simply be a consequence of the development of a drug seeking habit, since it was not seen in the FI15 control rats in which drug seeking was also habitual, but never under the control of cocaine-associated conditioned reinforcement. Instead, it emerged specifically in SOR rats in which cocaine-seeking habits had developed under, and were maintained by, the conditioned reinforcing properties of response-produced cocaine-associated CSs.

We have previously suggested that under these CS-controlled drug seeking conditions specific adaptations within amygdalo-striatal systems support associative representations we termed incentive habits (*21, 52, 54*) in which the response acquires the motivational value of the CS when it acts as a CRf. Here we demonstrate that these representations promote a flexibility in the goal and also the relative dominance by aDLS habit and pDMS goal-directed circuitries in the control, of an instrumental seeking behaviour characterized by the inflexibly of its initiation.

This study emphasizes the complex and dynamic nature of the behavioural factors underlying relapse in individuals with a long like history cocaine-seeking under the influence of Pavlovian drug CSs acting as conditioned reinforcers. In particular it brings into focus the potential role of negative urgency (*59*) and associated negative reinforcement (*61*) in the mechanisms by which maladaptive incentive drug seeking habits contribute to the chronicity of drug addiction (*10*) by increasing the likelihood and severity of drug seeking at relapse.

## Funding

This work was supported by a Programme Grant from the Medical Research Council to BJE and DB (MR/N02530X/1) and a grant from the Leverhulme Trust to DB (RPG-2016-117).

## Author contributions

DB, BJE, YPO and MF designed the experiments. MF, YPO, MP and CG performed all the behavioural experiments and the associated analyses. MF and YPO performed the histological assessments. YPO and NL performed the in-situ hybridization assays. MF produced all the figures. MF, YPO, BJE and DB wrote the manuscript.

## Competing interests

The authors have no financial disclosure or conflict of interest to report.

## Supplementary materials

Materials and methods

Figures S1-S6

References (1-13)

## Notes

### Competing Interest Statement

The authors have declared no competing interest.

