## Supplemental methods and figures for "Maladaptive cue-controlled cocaine-seeking habits promote increased relapse severity in rats"

This PDF file includes:

Materials and Methods

Figs. S1 to S6

Supplementary References (1-13)

### Materials and Methods

#### Subjects

Male Lister Hooded rats (n=120) (Charles River Laboratories, Kent, UK) weighing approximately 260-300g upon arrival were housed 2 per cage for a week of habituation. Rats had ad libitum access to standard chow and water and were maintained under a reversed 12-hr light/dark cycle (light ON between 7.00pm and 7.00am) at a constant temperature (21 ± 1° C). Rats were then food restricted (15-20g chow daily) and subjected to experimental procedures that were conducted 7 days a week. Procedures were conducted under the project license 70/8072 held by David Belin in accordance with the United Kingdom Animals (Scientific Procedures) Act 1986, amendment regulations 2012 following ethical review by the University of Cambridge Animal Welfare and Ethical Review Body (AWERB).

#### Drugs

Cocaine hydrochloride (Macfarlan-Smith, Edinburgh, UK and NIDA Drug Supply Programme) was dissolved in sterile physiological saline (0.9% sodium chloride) at a final concentration of 2.5g/L and stored at 4ºC. α-Flupenthixol (Sigma-Aldrich, Poole, UK) was dissolved in double-distilled water at a final concentration of 20mg/mL and stored at -20ºC.

#### Apparatus

Cocaine self-administration (SA) was conducted in standard operant conditioning chambers (31.8cm x 25.4cm x 26.7cm, Med Associates, St. Albans, VT, USA) located within a ventilated sound-attenuating cubicle as previously described (*1-3*). Each chamber was equipped with two retractable levers (4cm wide, 12cm apart, and 8cm from the grid floor), located on the front panel such as rats had access to both an active and an inactive lever, the location of which was randomised between individuals. Each test chamber was illuminated by one 3W light bulb (house light) during the experimental session and cue lights (2.5W) were located above each lever. A flexible tube, protected by a metal spring set up on a pivoting arm, was linked to a perfusion pump at one end and to the catheter on the other side. Operant chambers were controlled by MedPC (Med Associates, St. Albans, VT, USA) or Whisker software (*4*). All the SA experiments were performed 7 days a week during the dark phase of the light/dark cycle

#### Surgeries

##### *Intravenous Surgeries*

Rats were anaesthetised either by intramuscular injection of a ketamine/xylazine mixture in the experiments 1 and 2 (ketamine hydrochloride 90 mg/kg, Ketaset; xylazine 6.7 mg/kg, Rompun) or by isoflurane inhalation in the experiment 3 ($O_2$: 2L/min; 5% for induction and 2-3% for maintenance) and a silastic catheter (Camcaths, Ely, UK) was implanted into their right jugular vein as previously described (*1*). Rats were treated daily with oral antibiotic (Baytril 2.5%, 10mg/kg, Bayer) for one week from the day prior to surgery and with an analgesic agent (Metacam, 1mg/kg, Boehringer Ingelheim, UK) administered subcutaneously immediately prior to surgery and orally (1mg/kg daily) for the 3 consecutive days. Catheters were flushed daily with 0.1-0.2 ml of sterile physiological saline (0.9% sodium chloride) supplemented with heparin (50 IU/ml; Wockhardt UK) to maintain patency.

##### *Intracranial cannulations*

Stereotaxic surgeries were conducted under isoflurane anaesthesia ($O_2$: 2L/min, 5% isoflurane for induction and 2-3 % for maintenance) as previously described (*5*). Rats were positioned on the stereotaxic frame (David Kopf Instruments, Tujunga, CA, USA) and were implanted bilaterally with 22-gauge guide cannulae (Plastics One, Roanoke, VA, USA) positioned to lie 2mm above either the aDLS [anteroposterior (AP) +1.2, mediolateral (ML) ± 3, dorsoventral (DV) -3] or the pDMS [AP -0.4, ML ± 2.6 and DV -2.5] (*1, 6, 7*). AP and ML coordinates were measured from bregma, DV coordinates from the skull surface, with the incisor bar set at -3.3 mm. Cannulae were held in place using dental acrylic anchored to stainless steel crews tapped into the frontal and parietal bones of the skull. Obturators (PlasticsOne) were placed in the cannulae to maintain patency. Rats were treated with an analgesic agent (Metacam, 1mg/kg) administered subcutaneously immediately prior to surgery and orally (1mg/kg daily) for the 3 consecutive days.

#### Cocaine self-administration

Rats were trained to self-administer cocaine a week following intravenous surgery under a Fixed Ratio 1 (FR1) schedule of reinforcement for 6-8 days. Under this schedule of reinforcement, each active lever press resulted in the immediate delivery of cocaine (0.25mg/0.1 ml/5.7s/infusion) and a 20s presentation of the stimulus light located above the active lever (that will become the conditioned stimulus (Cs)). During these 20 seconds (time-out period) the house light was turned off and the levers were retracted. At the end of the time-out, the house light was turned back on, the CS switched off and the two levers inserted back into the chamber. This sequence of events associated with each cocaine infusion remained the exact same one throughout the self-administration (SA) procedure, regardless of the schedule of reinforcement subsequently used. Over the FR1 sessions, the number of drug infusions was limited to 30 over each 2-hour daily session. Inactive lever presses had no programmed consequences but were recorded to monitor the specificity of the instrumental response.

Rats were then trained to seek the drug under Fixed-interval (FI) schedules of reinforcement of increasing duration from 1 min (FI-1) to 15 min (FI-15) over 6 sessions. Rats were subjected to three sessions of FI-15 wherein drug infusion was available by pressing the active lever once a 15-min interval had elapsed. These sessions lasted until rats obtained 5 cocaine infusions or 2 hours had elapsed. After these 3 sessions, rats were either trained under a FI-15 schedule of reinforcement for ~15 days or trained to seek cocaine under the conditioned reinforcing properties of the drug-paired cue through a second-order schedule of reinforcement (FI15(FR10:S)), in which drug seeking, measured over prolonged periods of time (15 min) depended on the contingent presentation of the drug-paired CS (1s) every tenth active lever press. These CSs act as conditioned reinforcers, bridging delays to reinforcement and eventually facilitating the development of incentive habits (*6*). The number of infusions (which were available once the animal pressed ten time on the active lever after each 15-min interval has elapsed), was limited to 5 during each session (2 hours cut off).

##### *Intracranial infusions*

Vehicle or α-Flupenthixol (0.5ul/side/90s) was infused as previously described (*1, 6, 7*) by a syringe pump (Harvard Apparatus, Holliston, MA, USA) through 28-gauge steel hypodermic injectors (Plastics One) which protruded 2mm ventrally to the end of the guide cannula. Following each infusion, the injectors were maintained to allow the diffusion of the solution and test sessions began 5 min later. Rats were habituated to these infusions and associated manipulations at least 3 times prior to the first test session.

#### Histological assessment of cannula placements

At the end of the Experiments, Rats were anesthetised with a lethal dose of sodium pentobarbital (2 ml per rat, i.p., Euthatal, 200mg/ml; Genus Express) and perfused transcardially with 0.01M phosphate buffered saline (PBS) followed by 4% paraformaldehyde. The brains were harvested and post-fixed in 4% paraformaldehyde. Brains were then transferred to a 20% sucrose in 0.2M phosphate buffer solution and left to sit overnight. 60µm coronal sections were cut with a cryostat and stained with Cresyl Violet. Cannula placements were assessed under a light microscope and mapped onto standardised coronal sections of a rat brain stereotaxic atlas (**Fig. S5**) (*8*).

#### *In situ* hybridisation

After sacrifice, brains were immediately harvested and flash-frozen by immersion in -30/40ºC isopentane for 5 min and subsequently stored at -80ºC. Brains were processed into 12µm-thick coronal sections that were collected on gelatine-coated slides and stored at -80°C until further use. The previously described *in-situ* hybridization procedure (*9*) was carried out with oligonucleotide probe specifically complementary to the sequence of the mRNA of the immediate early gene C-Fos (nucleotides 159-203 of the NCBI Reference Sequence NM_022197.2) tailed by 3'OH incorporation of $^{35}$S-dATP (Perkin Elmer, UK) (1250 mCi/mmol) with a specificity of 2.5 x $10^6$ cpm/ml. Chiefly, following fixation and pre-hybridization treatments aiming at reducing non-specific hybridisation, slides were incubated overnight at 42ºC in the hybridisation buffer [*50% deionised formamide, 10%*

*dextran sulfate, 50ng/ml denaturated salmon sperm DNA, 5% Sarcosyl, 0.2% SDS, 1mM EDTA, 300nM NaCl, 5X Denhardt's in 2X standard sodium citrate (SSC)*] with the probe diluted at a concentration of 6.25ng/ml. Slides were then washed in decreasing concentration of SSC and dehydrated in increased ethanol concentration baths. Sections were exposed to Kodak Biomax MR films for 4 weeks at room temperature. Each film contained all the sections of a specific brain area for each of the rats tested. Films were revealed in a dark room. Pictures of each brain section were taken on a Northern light (Imaging Res Inc) light table with a Qicam (QImaging) camera equipped with a SIGMA 50 mm 1:2.8 DG MacroD Fast 1394 (Nikon) objective and subsequently analysed with ImageJ software. A region of interest was drawn for each striatal territory in which the optical density representative of the mRNA level was measured (according to the rat brain atlas (*8*)). The optical density in an mRNA-free part of the brain (i.e. a fibre tract) was defined as background and this value was subtracted to that obtained from the area of interest to compute the relative optical density used as the dependent variable in subsequent analyses.

#### Data and statistical analyses

Data are presented as box plots [median ±25% (percentiles) and Min/Max as whiskers] or mean ± 2SD and were analysed with Statistica Software 10 (Statsoft) or Statistical Package for Social Sciences (SPSS, 21.0). Assumption for parametric analyses, namely homogeneity of variance, sphericity and normality of distribution were verified prior to each analysis with Cochran, Mauchly and Shapiro-Wilk's tests respectively.

Lever presses during baseline, at relapse and during acquisition of both cocaine SA and cocaine seeking across the fixed interval schedules of reinforcement were analysed using repeated-measures analysis of variance (ANOVAs) with lever (active and inactive) and session as within-subject factors, and group (FI15, SO, ShA,LgA…), experimental condition of abstinence (instrumental deprivation with or without non-contingent cocaine infusions), localisation of α-flupenthixol (aDLS/pDMS) and experimental conditions at test (CS+/CS-) as between-subject factors. Because of between-session variability in performance, data pertaining to the acquisition of cue-controlled cocaine seeking behaviour were analysed across 3-days blocks (including a first block of 3 FI15 sessions).

The confirmation of significant main effects and differences among individual means were analysed using the Newman-Keuls post-hoc test. For all analyses, significance was set at $\alpha$= 0.05 and effect sizes were reported as partial eta squared ($\eta_p^2$).

#### Specific experimental procedures

*Experiment 1: Characterisation of the reliance on the instrumental deprivation effect on pharmacological withdrawal.*

Following acquisition of cocaine SA (6 days) under FR1 and subsequent FI training, rats (n=47) were split in two groups with similar rate of responding. One group continued to seek cocaine under a FI15 schedule of reinforcement (FI15 group, n=23) while the other group was trained under a second order (SO) schedule of reinforcement (SO group, n=24). Rats were trained for 15 days under these differential conditions and their propensity to relapse to cocaine seeking was subsequently tested under different experimental conditions (experiments 1a, 1b, 1c and 1d).

To measure the effects of abstinence on drug seeking behaviour at relapse, rats with a history of 15 days of training under FI15 (n=11) or SO (n=12) underwent 3 days of imposed abstinence during which they were left undisturbed in their home-cages, handled only once daily for the routine flushing of the intravenous catheter. On the 4th day, rats returned to the SA chamber and relapse to cocaine seeking behaviour was measured over the 15 min drug-free interval. The magnitude of relapse to drug seeking was calculated across all experiments as the comparison between the active lever presses during the relapse session and the average of that expressed over the 15 min drug-free interval of the 2 preceding baseline sessions (experiment 1a). Then, rats underwent at least 3 baseline sessions until they recovered a stable drug seeking behaviour.

In order to test whether the increased drug seeking behaviour at relapse, called instrumental rebound, was driven by acute drug withdrawal, rats were placed each day over the 3-days abstinence period in the SA chamber for 2 hours, where they had no access to the levers and no lights were presented, and they received non-contingent cocaine infusions every 15 minutes, up to 5 infusions per day.

Importantly, rats were not given the opportunity instrumentally to respond. Rats from each group either received non-contingent cocaine infusions or were left undisturbed in the SA chamber (SO group, n=9 and n=12 respectively; FI15 group, n=10 and n=12 respectively). After these 3 days of abstinence during which they could not perform instrumental responses, i.e., instrumental deprivation, rats were placed back into their SA chambers and underwent a relapse test (experiment 1b). Then, rats were given at least 3 baseline sessions until they recovered a stable drug seeking behaviour.

To test whether the increased drug seeking behaviour at relapse displayed by the SO group following abstinence (Fig. 1) was attributable to an increased sensitivity to conditioned reinforcement, rats from the SO group (n=23) which, after 3 days of imposed abstinence, were returned to the SA chamber and given the opportunity to seek the drug with (CS+, n=12) or without (CS-, n=11) contingent presentation of the CS. (experiment 1c).

To investigate the neural correlates, c-Fos mRNA levels associated with baseline seeking behaviour or rebound at relapse were measured in the dorsal striatum using in situ hybridisation assays, rats from the SO group underwent at least 3 baseline sessions prior to be subjected to either abstinence for 3 days whereby rats were left undisturbed in their home cage (handled only once daily for the routine flushing of the intravenous catheters) or to three additional baseline sessions. Their relapse to drug seeking behaviour was subsequently tested under a drug-free 15-min seeking session (i.e., first interval of a SO schedule of reinforcement). Rats were immediately returned to their home cage and 45 min after were decapitated and their brains harvested and snap frozen in -30/-40ºC isopentane and stored at -80ºC. Due to technical constraints, i.e. the limited number of slides fitting in an X-Ray cassette, the in situ hybridisation assays were carried out on 8 rats from each condition (i.e. baseline and abstinence) (experiment 1d).

Over the entire experiment, three rats from the SO group and two from the FI15 group were excluded from analyses during the SA training and the experiments 1b and 1c due to loss of catheter patency.

*Experiment 2. Influence of previous cocaine exposure on the instrumental deprivation effect.*

After acquisition of cocaine SA over 8 daily 1-hour sessions under continuous reinforcement (FR-1), 12 rats were maintained under the same short access (ShA; 1h/day, 30 infusions max) conditions while the other 12 were given 6-hour daily (150 infusions max) extended access (LgA) to cocaine (*2, 10, 11*). After 12 sessions and the establishment of escalation of cocaine SA in LgA rats, both groups were trained to seek cocaine under incremental FI schedules of reinforcement and subsequently, under a SO schedule of reinforcement as described above.

After 15 sessions of SO schedule of reinforcement, the influence of a history of escalated SA on the rebound effect was measured as described above. Thus, rats were exposed to 3 days of imposed abstinence during which they were left undisturbed in their home-cages, handled only once daily for the routine flushing of the intravenous catheter. On the 4$^{th}$ day, rats returned to the SA chamber and their relapse to cocaine seeking behaviour was measured over the 15 min drug-free interval (**experiment 2a**). After at least 3 baseline sessions (until stable level of responding), rats underwent the same procedure but were tested at relapse in a CS- context as described above (**experiment 2b**). One rat from the ShA group and four from the LgA group were excluded from the analysis due to loss of catheter patency.

*Experiment 3. Causal identification of the striatal locus of control of the rebound.*

Having established by post-mortem assessments (see experiment 1c) that the increased cocaine seeking behaviour at relapse following abstinence was associated with an increased expression of C-Fos in the posterior dorsomedial striatum (pDMS), we sought causally to investigate the reliance of drug seeking, otherwise dependent on aDLS dopaminergic (DA) mechanisms (*1, 6, 7, 12*), on pDMS DA mechanisms at rebound. For this, we tested the sensitivity of baseline or post-abstinence seeking behaviour to DA receptors blockade either in the aDLS or the pDMS.

Two cohort of rats (n=24 each) were implanted with bilateral guide cannula targeting either the aDLS or the pDMS and, a week later, an indwelling catheter in their right jugular vein. At least 5 days later, rats were trained to acquire cocaine SA under continuous reinforcement for 5 daily sessions before being progressively trained to seek cocaine under FI schedules of reinforcement with increasing interval duration, as described above, prior to be subjected to 27 SA sessions under SO schedule of reinforcement. Then, we measured the effects of bilateral infusions of α-flupenthixol on seeking behaviour at relapse following 3 days of imposed abstinence. Rats received intracranial infusions of vehicle or α-flupenthixol (10μg/infusion) (dose based on previous studies from our laboratory (*1, 5-7*)) either at baseline or at the relapse test following a latin square design. The reliance of drug seeking at baseline or during the expression of rebound on the different dorsal striatal territories was calculated as the percentage of active lever presses when injected with α-flupenthixol over the active lever presses when injected with vehicle.

In order further to characterise the behavioural and psychological mechanisms of the instrumental deprivation effect, shown in the experiment 1 not to depend on pharmacological withdrawal, we tested the specificity of the effect on the inability to express seeking, but not taking responses. Rats (n=11) were given the opportunity to self-administer 1 cocaine infusion every 15 min without the opportunity to make their usual seeking responses for 3 days prior to being challenged in a relapse test. For this, after at least 3 baseline sessions following the last intracerebral challenge, rats were placed for 3 consecutive daily sessions in their SA chambers. The session began with the illumination of the house-light, but rats had no access to the levers. After an interval of 15 min had elapsed, both active and inactive levers were inserted in the chamber and 1 active lever press (a taking response) resulted in the immediate delivery of a cocaine infusion (0.25mg/0.1 ml/5.7s/infusion) and a 20s presentation of the stimulus light located above the active lever. During these 20 seconds (time-out period) the house light was turned off and the levers were retracted. At the end of the time-out, the house light was turned back on, the CS switched off, but the two levers remained retracted. This cycle and the opportunity for each individual to self-administer cocaine was repeated after another 15 min interval had elapsed until rats received 5 infusions (or 90 min cut off). Following these 3 sessions, rats were allowed to express their seeking behaviour during a relapse challenge session (with levers available from the onset of the session).

Rats were perfused and their brains harvested and subsequently processed for histological assessment of the injection sites. Due to loss of catheter patency or cannula misplacements, the final number of rats included in the analysis was eleven from the cohort having been implanted with cannula targeting the aDLS and ten from the cohort with cannula targeting the pDMS.

### Supplementary Figures

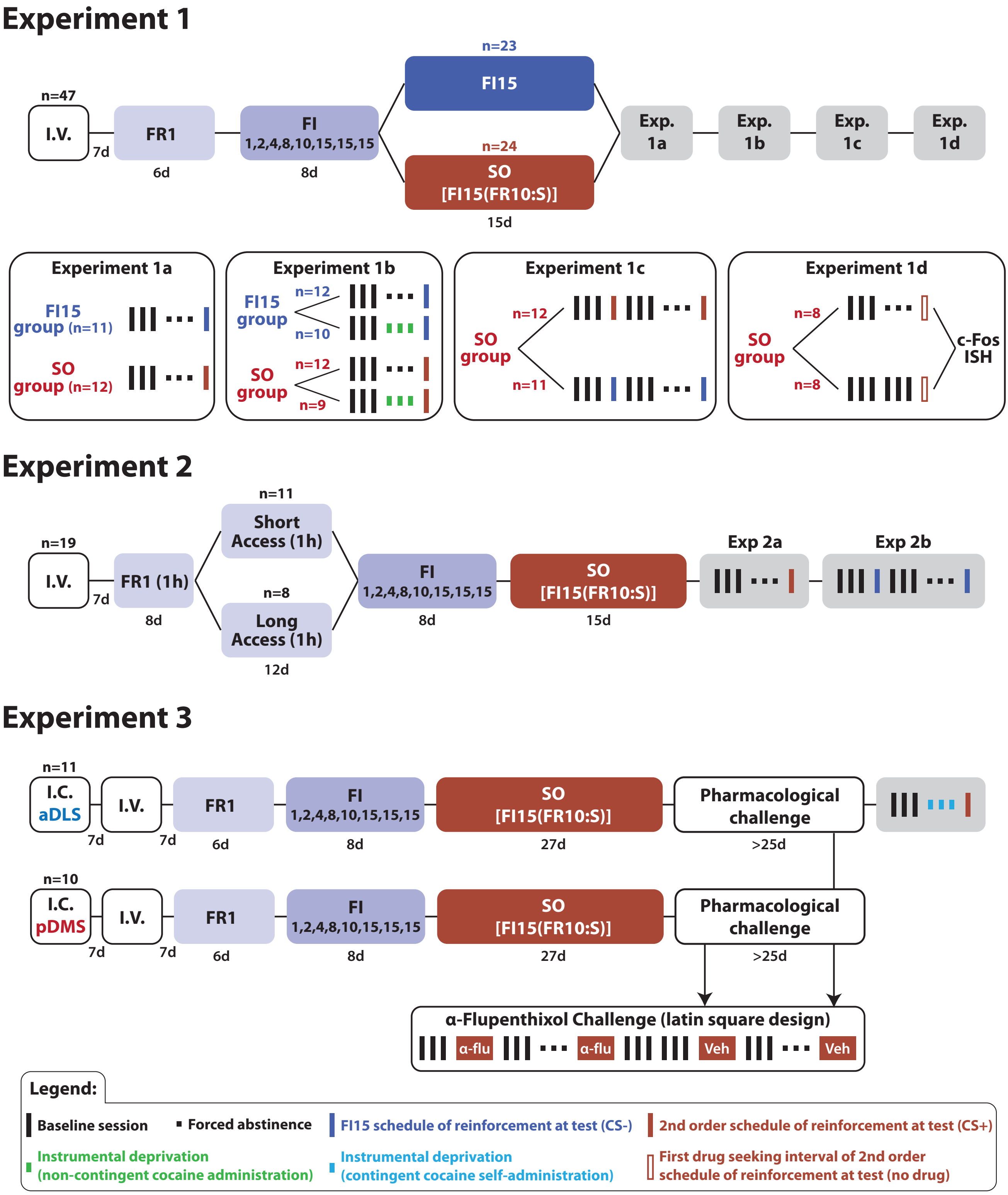


#### Figure S1: Experimental timeline of the experiments

**Experiment 1:** Following acquisition of cocaine SA (6 days) under FR1 and subsequent FI training, rats (n=47) were split in two groups displaying similar rate of responding. One group remained exposed to the same FI15 schedule of reinforcement (FI15 group, n=23) for 15 days while the other group was trained to seek cocaine under the control of the drug-paired cues acting as conditioned reinforcers, as operationalised under a second order (SO) schedule of reinforcement (SO group, n=24). After 15 days of training under those conditions, their propensity to relapse to cocaine seeking behaviour was subsequently challenged under different experimental conditions (experiments 1a, 1b, 1c and 1d). Experiment 1a: half the rats from each group (FI15, n=11 and SO, n=12) were subjected to 3 days of forced abstinence. On the 4$^{th}$ day, rats returned to the SA chamber and their propensity to relapse to cocaine seeking behaviour was measured over the 15 min drug-free interval. The magnitude of relapse to drug seeking was calculated across all experiments as the comparison between the active lever presses during the relapse session and the average of that expressed over the 15 min drug-free interval of the 2 preceding baseline sessions. Experiment 1b: The role of the pharmacological withdrawal (from cocaine) in the rebound at relapse was assessed in the entire cohort of 43 rats trained under SO (n=21) or FI15 (n=22) schedule of reinforcement. After at least 3 baseline sessions, rats were placed into the SA chamber for 2 hours once daily over 3 days. They had no access to the levers and no lights were presented at any moment. Rats from each group (SO group, n=9 and FI15 group, n=10) received a non-contingent cocaine infusion every 15 minutes (5 infusions per session), while the other rats (n=12 per group) received no infusion and remained undisturbed in the SA chamber for the same duration. After these three days, rats were placed back into their SA chambers where they were subjected to relapse test. Experiment 1c: The influence of the conditioned reinforcement on the instrumental deprivation effect shown by SO rats was assessed in a group of 23 individuals, which, after at least 3 baseline sessions and 3 days of forced abstinence were returned to the SA chamber and given the opportunity to seek the drug with (CS+, n=12) or without (CS-, n=11) contingent presentation of the CS. Experiment 1d: A subset of SO rats was used to identify the striatal correlates of rebound at relapse. Rats underwent at least 3 baseline sessions (until stable level of responding) prior to be subjected to either forced abstinence for 3 days (n=8) or to three additional baseline sessions (n=8). Their propensity to relapse to drug seeking behaviour was subsequently tested under a drug-free 15-min seeking session (i.e. first interval of a SO schedule of reinforcement). Rats were immediately returned to their home cage and 45 min after were decapitated and their brains harvested and snap frozen to carry out further *in situ* hybridisation assays. **Experiment 2:** After acquisition of cocaine SA over 8 1-hour daily sessions under continuous reinforcement (FR-1), 12 rats were maintained under the same short access (ShA; 1h/day, 30 infusions max) conditions while the other 12 were given 6-hour daily (150 infusions max) extended access (LgA) to cocaine (*2, 10, 11*). After 12 sessions and the establishment of escalation of cocaine SA in LgA rats, both groups were trained to seek cocaine under incremental FI schedules of reinforcement and subsequently, under the control of the drug-paired cues as operationalised under a SO schedule of reinforcement as described above. After 15 sessions under SO schedule of reinforcement, the influence of a history of escalated SA on the rebound at relapse was measured. Rats were exposed to 3 days of forced abstinence during which they were left undisturbed in their home-cages, On the 4$^{th}$ day, rats returned to the SA chamber and their propensity to relapse to cocaine seeking behaviour was measured over a 15 min drug-free interval. After at least 3 baseline sessions, rats were subjected to the same procedure but were tested at relapse in an CS- context as described above. **Experiment 3:** Two cohort of rats (n=24 each) were implanted with bilateral guide cannula targeting either the aDLS (n=11) or the pDMS (n=10) and, a week later, an indwelling catheter in their right jugular vein. At least 5 days later, rats were trained to acquire cocaine SA under continuous reinforcement for 5 daily sessions before being progressively trained to seek cocaine under FI schedules of reinforcement with increasing interval duration, as described above, prior to be subjected to 27 SA sessions under SO schedule of reinforcement. Then, we measured the effects of bilateral infusions of α-flupenthixol on seeking behaviour at relapse following 3 days of forced abstinence. Rats received intracranial infusions of vehicle or α-flupenthixol (10μg/infusion) either during baseline or at relapse test following a latin square design. In order further to characterise the behavioural and psychological mechanisms of the instrumental deprivation effect, shown in the experiment 1 to not be dependent on pharmacological withdrawal, we tested the specificity of the effect to the inability to express seeking, but not taking responses. Thus, rats (n=11) were given the opportunity to self-administer 1 cocaine infusion every 15 min without the opportunity to express seeking responses for 3 days prior to be challenged in a relapse test. For this, after at least 3 baseline sessions following the last intracerebral challenge, rats were placed for 3 consecutive daily sessions in their SA chambers. The session started with the illumination of the house-light, but rats had not access to the levers. After an interval of 15 min had elapsed, both active and inactive levers were inserted in the chamber and 1 active lever press resulted in the immediate delivery of a cocaine infusion and a 20s presentation of the stimulus light located above the active lever. During these 20 seconds the house light was turned off and the levers were retracted. At the end of the time-out, the house light was turned back on, the CS switched off, but the two levers remained retracted. This cycle and the opportunity for each individual to self-administer cocaine was repeated after another 15 min interval had elapsed until rats received 5 infusions (or 90 min cut off). Following these 3 sessions, rats were allowed to express their seeking behaviour during a relapse challenge session (with levers available from the onset of the session). Rats were then perfused, and their brains harvested and subsequently processed for histological assessment of the injection sites.

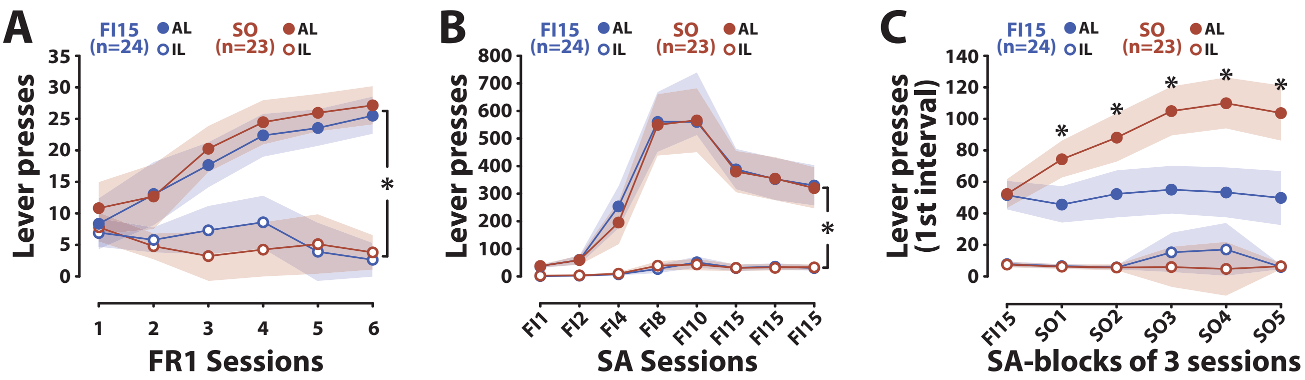


#### Figure S2: Acquisition and maintenance of drug seeking behaviour of rats trained under SO and FI15 schedules of reinforcement in the experiment 1.

**A)** Rats later allocated to the FI-15 (n=24) or SO (n=23) group quickly learned instrumentally to respond for IV cocaine infusions as shown by the rapid development of a discrimination between the active (AL) and inactive (IL) lever presses over 6 daily sessions under continuous reinforcement [main effect of lever: $F_{1,45}$=113.95, p≤.001, $\eta_p^2$=.72; session: $F_{5,225}$=17.42, p≤.001, $\eta_p^2$=.28; group: $F_{1,45}$<1; lever x session interaction: $F_{5,225}$=36.88, p≤.001, $\eta_p^2$=.45; lever x group interaction: $F_{1,45}$=1.14, p>.05 and lever x session x group interaction: $F_{5,225}$=1.27, p>.05]. **B)** Rats progressively learnt to seek cocaine interval of time of increasing duration, from 1 min (FI1) to 15 min (FI15) [main effect of lever: $F_{1,45}$=233.87, p≤.001, $\eta_p^2$=.84; session: $F_{7,315}$=114.39, p≤.001, $\eta_p^2$=.72; group: $F_{1,45}$<1; lever x session interaction: $F_{7,315}$=99.57, p≤.001, $\eta_p^2$=.69; lever x group interaction: $F_{1,45}$<1 and lever x session x group interaction: $F_{7,315}$<1]. **C)** The introduction of the response-produced CSs in the SO group resulted in the well characterised invigoration of instrumental seeking responses [main effect of lever: $F_{1,45}$=252.63, p≤.001, $\eta_p^2$=.85; block of sessions: $F_{5,225}$=9.99, p≤.001, $\eta_p^2$=.18; group: $F_{1,45}$=11.51, p≤0.01, $\eta_p^2$=.20; lever x block of sessions interaction: $F_{5,225}$=6.59, p≤.001, $\eta_p^2$=.13; lever x group interaction: $F_{1,45}$=27.83, p≤.001, $\eta_p^2$=.38 and lever x block of sessions x group interaction: $F_{5,225}$=8.66, p≤.001, $\eta_p^2$=.16]. *p≤.05

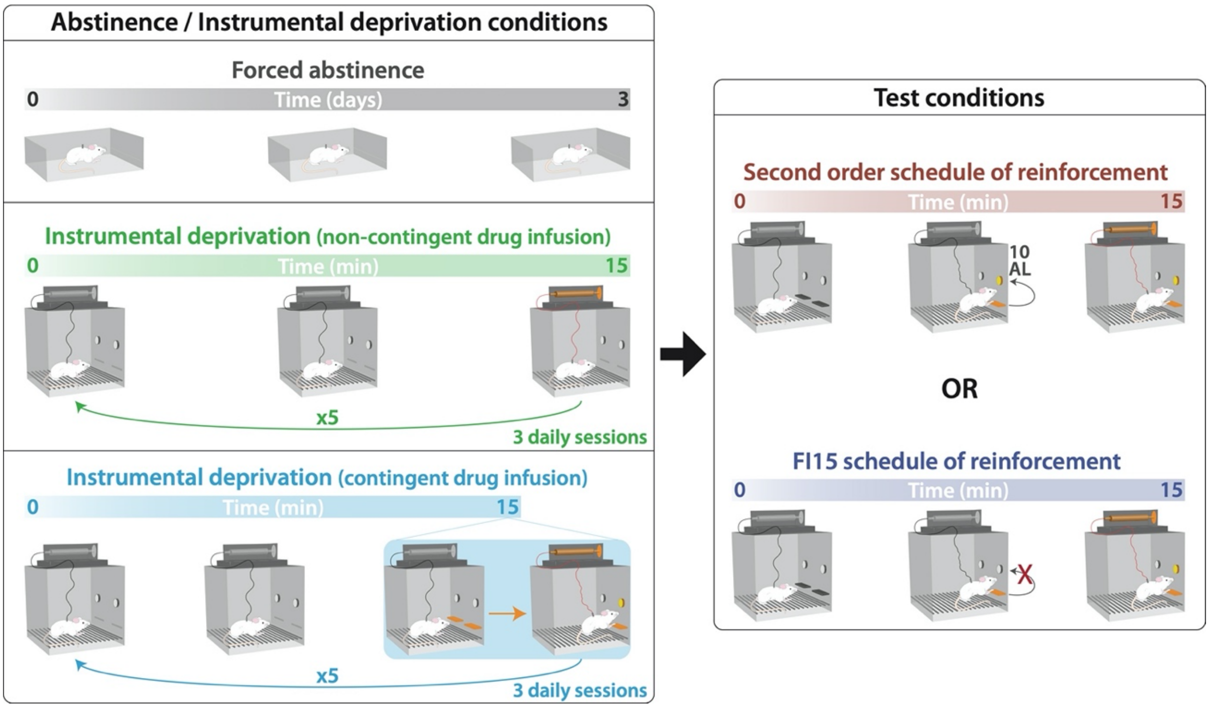


**Figure S3: Schematic representation of the abstinence, instrumental deprivation and test conditions.**

**Left panel-** Rats were subjected to one of the three types of behavioural manipulations prior to the subsequent assessment of their tendency to relapse under different test conditions. **Top-** Rats were subjected to 3 days of forced abstinence during which they were left undisturbed in their home cages while their catheters were flushed daily with sterile heparinized saline. **Middle-** Rats were placed into the SA chamber for 75 minutes once daily for the same 3-day period. They had no access to the levers and no lights were presented at any moment. Rats received a non-contingent cocaine infusion every 15 minutes (5 infusions per session), as they otherwise received, albeit contingently, during baseline sessions. **Bottom-** Rats were placed for 3 consecutive daily sessions in their SA chambers with no access to the levers. Each day, after an interval of 15 min had elapsed, both active and inactive levers were inserted in the chamber and 1 active lever press resulted in the immediate delivery of a cocaine infusion associated with the previously described 20s time-out period. At the end of the time-out, the two levers remained retracted preventing the expression of seeking behaviour. This cycle and the opportunity for each individual to self-administer cocaine was repeated after another 15 min interval had elapsed until rats received 5 infusions. **Right panel-** Rats' propensity to relapse following either abstinence or instrumental deprivation was assessed using one of the two following schedules of reinforcement. **Top-** Rats were given the opportunity to seek cocaine under the conditioned reinforcing properties of the drug-paired cue as in a second-order schedule of reinforcement (SOR: FI15(FR10:S)), in which drug seeking, measured over 15 min was invigorated by the contingent presentation of the drug-paired CS (1s) every tenth active lever press. Drug infusion was available once the animal had pressed ten times on the active lever after each 15-min interval has elapsed. **Bottom-** Rats were given the opportunity to seek cocaine under a FI15 schedule of reinforcement in which they had access to both inactive and active levers throughout and received a drug infusions upon pressing the active lever after each 15 min interval had elapsed.

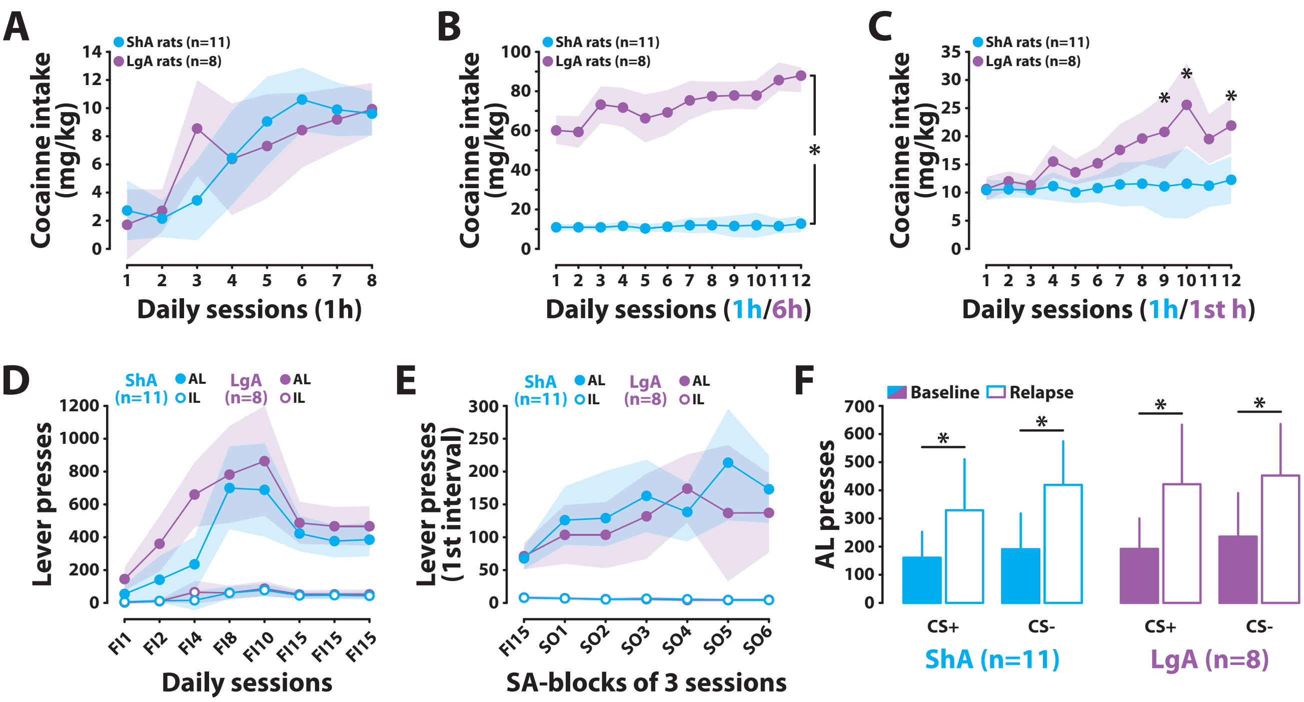


**Figure S4: Differential access to cocaine self-administration did not influence the propensity to acquire cocaine seeking, its potentiation by the conditioned reinforcing properties of the drug-paired cue and the rebound at relapse after instrumental deprivation.**

**A)** Rats later allocated to the Short Access (ShA, n=11) or Long Access (LgA, n=8) group, acquired cocaine SA similarly over 8 1-hour daily sessions under continuous reinforcement (the deviation on day 3 was due to 1 individual). [main effect of group: $F_{1,17}<1$; session: $F_{7,119}=174.43$, $p\leq.001$, $\eta_p^2=.50$ and group x session interaction: $F_{7,119}=2.30$, $p\leq.05$, $\eta_p^2=.03$, Left panel]. **B)** Introduction of 6-hour extended daily access in the LgA group resulted in a robust escalation of intake as compared to the intake shown by the ShA group that remained stable over 12 sessions [main effect of group: $F_{1,17}=272.73$, $p\leq.001$, $\eta_p^2=.94$; session: $F_{11,187}=5.13$, $p\leq.001$, $\eta_p^2=.23$ and group x session interaction: $F_{11,187}=4.18$, $p\leq.05$, $\eta_p^2=.20$, Right panel]. **C)** The escalation of cocaine intake shown by LgA rats was also apparent during the first hour of each daily session, as previously reported (*11, 13*) [main effect of group: $F_{1,17}=11.62$, $p\leq.01$, $\eta_p^2=.40$; session: $F_{11,187}=6.31$, $p\leq.001$, $\eta_p^2=.27$ and group x session interaction: $F_{11,187}=4.13$, $p\leq.001$, $\eta_p^2=.20$]. **D)** Both group similarly learnt progressively to seek cocaine interval of time of increasing duration, from 1 min (FI1) to 15 min (FI15) [main effect of lever: $F_{1,17}=103.57$, $p\leq.001$, $\eta_p^2=.86$; session: $F_{7,119}=24.57$, $p\leq.001$, $\eta_p^2=.59$; group: $F_{1,17}=3.08$, $p>.05$; lever x session interaction: $F_{7,119}=18.12$, $p\leq.001$, $\eta_p^2=.52$; lever x group interaction: $F_{7,119}=3.24$, $p>.05$ and lever x session x group interaction: $F_{7,119}=1.31$, $p>.05$]. **E)** The differential access to cocaine SA did not influence the acquisition of cue-controlled drug seeking behaviour. Indeed, introduction of response-produced CSs under a second order schedule of reinforcement resulted in a similar invigoration of instrumental seeking responses in ShA and LgA rats [main effect of lever: $F_{1,17}=71.91$, $p\leq.001$, $\eta_p^2=.81$; block of sessions: $F_{6,102}=7.27$, $p\leq.001$, $\eta_p^2=.30$; group: $F_{1,17}<1$; lever x block of sessions interaction: $F_{6,102}=8.49$, $p\leq.001$, $\eta_p^2=.33$; lever x group interaction: $F_{1,17}<1$ and lever x block of sessions x group interaction: $F_{6,102}=1.95$, $p>.05$]. **F)** Similarly, the magnitude of the instrumental deprivation effect, as measured as an increased instrumental responding after 3 days of forced abstinence, was the same in LgA and ShA rats [main effect of lever: $F_{1,17}=94.44$, $p<.001$, $\eta_p^2=.85$; abstinence: $F_{1,17}=11.68$, $p<.01$, $\eta_p^2=.41$; and lever x abstinence interaction: $F_{1,17}=11.42$, $p<0.01$, $\eta_p^2=.40$]. The instrumental deprivation effect was not dependent either on the contingent presentation of the conditioned reinforcer at relapse [no main effect of CS presentation, of group and no (group x

CS x lever); (group x CS x abstinence) and (group x CS x abstinence x lever) interaction: all Fs<1]. *$p \leq .05$

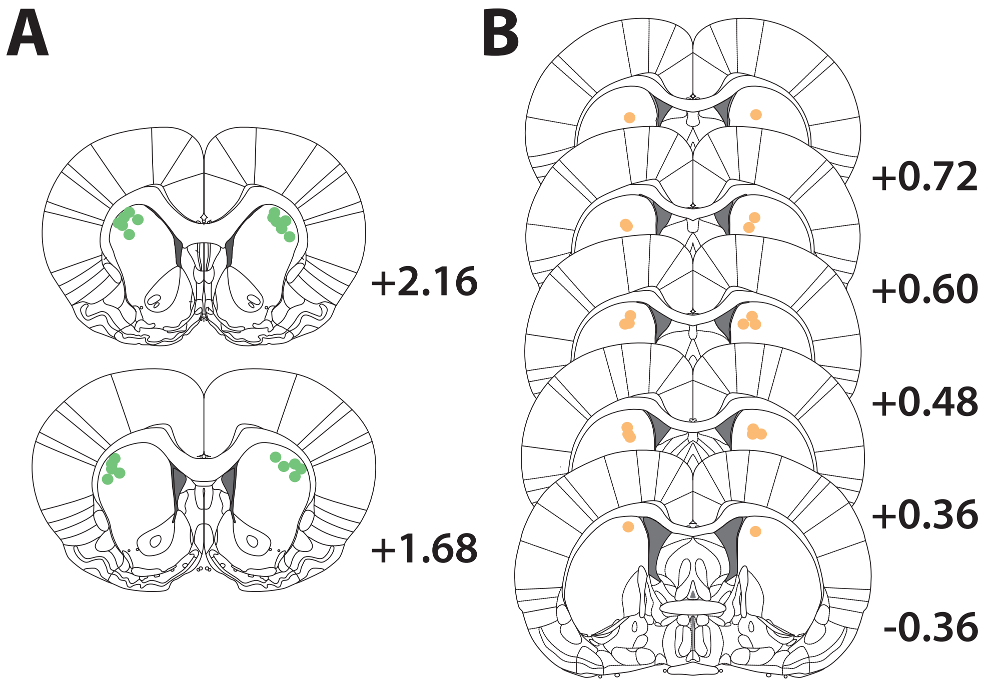


**Figure S5: Schematic representation of the injection sites targeting either the aDLS or the pDMS.**

Following sectioning of the perfused brain and staining of the sections with Cresyl Violet, the sites of injections were revealed under a light microscope to be located in the aDLS between +2.16 and +1.68 in the anteroposterior axis from bregma **(A)** or in the pDMS between +0.72 and -0.36 from bregma **(B)** according to the rat brain atlas (*8*).

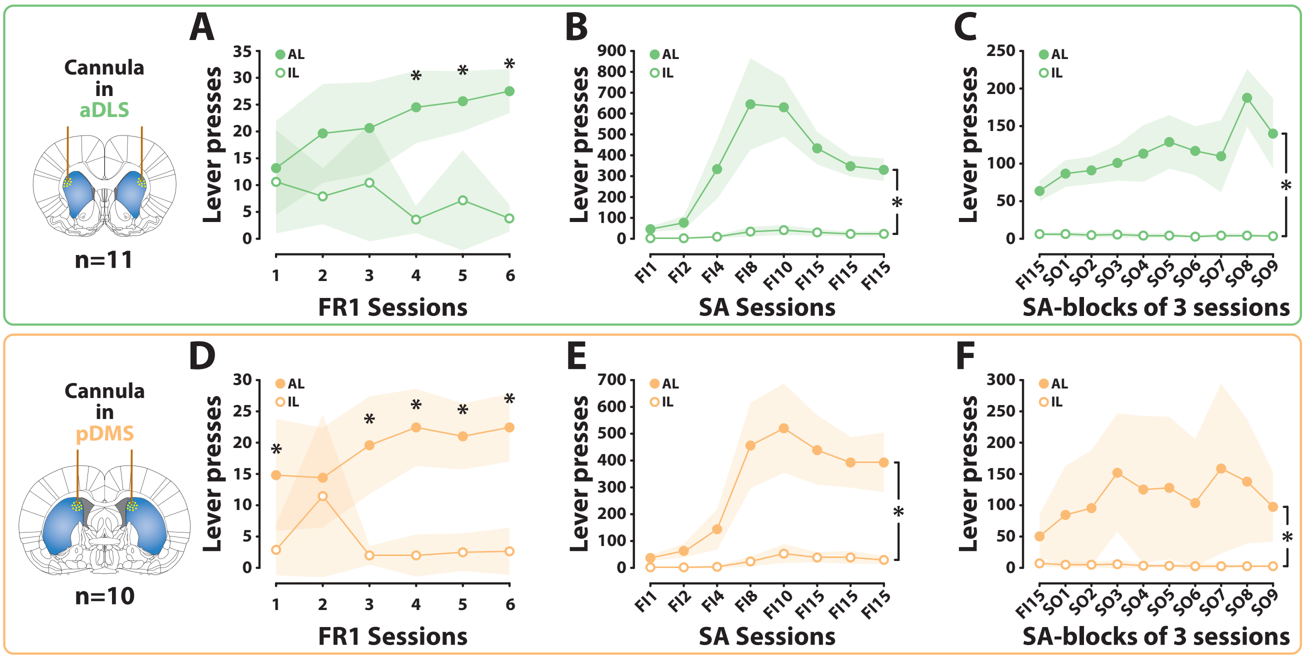


**Figure S6: Rats in which the reliance of instrumental seeking on dopaminergic mechanisms in the aDLS or pDMS was tested acquired and maintained cue-controlled cocaine seeking behaviour similarly.**

**A and D)** Rats (n=21) that underwent intracranial surgeries for the bilateral implantation of cannulas targeting either the aDLS (n=11) or pDMS (n=10) quickly learned instrumentally to respond for IV cocaine infusions as shown by the rapid development of a discrimination between the active (AL) and inactive (IL) lever presses over 6 daily sessions under continuous reinforcement [aDLS group: main effect of lever: $F_{1,10}$=27.26, p<.001, $\eta_p^2$=.73; session: $F_{5,50}$<1 and lever x session interaction: $F_{5,50}$=6.48, p<.001, $\eta_p^2$=.39; pDMS group: main effect of lever: $F_{1,9}$=21.50, p<.001, $\eta_p^2$=.70; session: $F_{5,45}$=1.41, p>.05 and lever x session interaction: $F_{5,45}$=3.58, p<001, $\eta_p^2$=.28]. **B and E)** Following acquisition of cocaine SA under continuous reinforcement, aDLS and pDMS rats acquired readily to respond for cocaine over intervals of time of increasing duration from 1 min (FI1) to 15 min (FI15) [aDLS group: main effect of lever: $F_{1,10}$=137.68, p<.001, $\eta_p^2$=.93; session: $F_{7,70}$=28.03, p<.001, $\eta_p^2$=.74 and lever x session interaction: $F_{7,70}$=22.70, p<.001, $\eta_p^2$=.69; pDMS group: main effect of lever: $F_{1,9}$=74.60, p<.001, $\eta_p^2$=.89; session: $F_{7,63}$=25.64, p<.001, $\eta_p^2$=.74 and lever x session interaction: $F_{7,63}$=29.30, p<001, $\eta_p^2$=.76]. **C and F)** The introduction of response-produced CSs under a second order schedule of reinforcement resulted in an invigoration of instrumental seeking responses in both group [aDLS group: main effect of lever: $F_{1,10}$=164.56, p<.001, $\eta_p^2$=.94; block of sessions: $F_{9,90}$=6.73, p<.001, $\eta_p^2$=.40 and lever x block of session interaction: $F_{9,90}$=7.33, p<001, $\eta_p^2$=.42; pDMS group: main effect of lever: $F_{1,9}$=40.36, p<.001, $\eta_p^2$=.81; block of sessions: $F_{9,81}$=2.82, p<.01, $\eta_p^2$=.24 and lever x block of sessions interaction: $F_{9,81}$=3.18, p<.01, $\eta_p^2$=.26]. *p≤.05
